# Evaluating the performance of photon- and electron-based fragmentation methods in Omnitrap-LCMS analysis of N-glycopeptides

**DOI:** 10.64898/2025.12.10.693381

**Authors:** Nikita Levin, Shabaz Mohammed

**Affiliations:** Rosalind Franklin Institute, Harwell Campus, OX11 0QX Didcot, U.K.; Department of Pharmacology, University of Oxford, OX1 3QT Oxford, U.K.; Department of Biochemistry, University of Oxford, OX1 3QU Oxford, U.K.; Department of Chemistry, University of Oxford, OX1 3TA Oxford, U.K.

**Keywords:** Glycoproteomics, mass spectrometry, ultraviolet photodissociation, electron ionization dissociation

## Abstract

To date, collision-induced dissociation and methods based on electron transfer dissociation are considered the standard approaches for the mass spectrometry analysis of N- and O-glycopeptides, respectively, allowing for identification of both peptide and glycan compositions. In recent years, alternative fragmentation methods such as ultraviolet photodissociation (UVPD) and more energetic versions of electron-based techniques (such as electron ionisation dissociation, EID) have been shown to be useful for the analysis of glycopeptides, producing rich information on the glycopeptide structure, including types of glycosidic linkages. We evaluated ultraviolet photodissociation (UVPD), electron ionization dissociation (EID), electron capture dissociation (ECD), and activated-ion ECD (AI-ECD) using an Orbitrap–Omnitrap hybrid for LC–MS analysis of complex N-glycopeptides. Both UVPD and EID generated extensive peptide, glycosidic, and cross-ring fragments, enabling detailed structural characterization. While ECD alone produced few glycopeptide identifications, AI-ECD significantly improved yields through supplemental vibrational activation. UVPD and EID achieved comparable identification efficiencies to stepped collisional dissociation and provided additional linkage information. These results establish the Omnitrap as a powerful platform for comprehensive glycoproteomic analysis and highlight the need for enhanced computational tools to interpret complex UVPD and EID spectra.

## Introduction

Protein glycosylation is a covalent attachment of an (oligo)saccharide to specific amino acid residues within the polypeptide chain. Glycosylation is one of the most abundant and complex posttranslational modifications (PTMs) and plays a crucial role in multiple biological processes ranging from cell signalling and immune response to protein folding and protein stability.^1, 2^ Glycoproteins and glycopeptides are often characterized by various degrees of glycosylation site occupancy (macroheterogeneity) and high diversity of glycan structure (microheterogeneity) including glycan composition, branching, and types of glycosidic bonds.^3, 4^ This intricate nature of glycopeptides poses a significant challenge to the analysis of complex glycopeptide mixtures by conventional analytical techniques such as reverse phase liquid chromatography coupled with mass spectrometry (RPLC-MS).^5-7^ In particular, collision induced dissociation (CID) often fails to sufficiently characterize both peptide and glycan structures and firmly identify the glycosylation site due to high acid lability of glycosidic ether bonds resulting in their preferential dissociation in a typical CID experiment.^6^ To address this shortcoming, a variety of alternative fragmentation techniques have been suggested, that include methods based on the use of electrons of various energies,^8-19^ infrared radiation,^10, 11, 20^ ultraviolet photodissociation (UVPD),^21-23^ and combinations thereof.^24^ While electrons with relatively low energies (<5 eV) provide peptide fragments complementary to CID,^9-12, 14, 15^ high-energy electrons and UV light generate all types of peptide main-series fragments, dissociate glycosidic bonds and produce cross-ring fragments of sugars.^16-18, 21, 23, 25, 26^ These fragments, when properly annotated, allow to confidently confirm the glycopeptide composition and provide valuable information about the types of glycosidic linkages between saccharides which may be useful for unravelling their biological function.^27-29^ This level of information potentially enables direct analysis of glycosidic bonds by means of MS/MS sequencing of glycopeptides without chemical derivatization of saccharides, which is currently the analytical method of choice.^30^

Despite ever-increasing attention from the glycopeptide community towards UVPD and electron-based dissociation techniques (ExD), literature providing fragmentation insight and utilisation approaches, especially on the LCMS scale, is sparce, not least because of prohibitive requirements to instrumentation, (subsequent) lack of mechanistic understanding, and insufficient tools for data analysis. To address some of these issues, we have previously employed a hybrid Orbitrap-Omnitrap platform^31^ to characterize UVPD, EID and ECD in a series of direct-infusion experiments of a few standard polypeptides^32^ and LCMS analyses of a variety of highly complex peptide mixtures.^33^ In this paper, we extend our characterisation of UVPD, EID, ECD, and activated-ion ECD (AI-ECD) to LC-MS analyses of complex N-glycopeptide mixtures.

## Methods

### Materials

Two standard glycopeptides with amino acid sequence KVANKT and glycoforms A2G2S2 (HexNAc4Hex5NeuAc2) and FA2 (HexNAc4Hex3Fuc) were purchased from Ludger Ltd, see Supplementary Figure S1d for detailed annotation of their structures. Human serum, solvents, and chemicals for sample preparation were obtained from Sigma Aldrich. Trypsin was supplied by Promega, and LysC was purchased from Fujifilm Wako. PNGase F kit containing PNGase F powder and GlycoBuffer 2 (50 mM Sodium Phosphate) was acquired from New England Biolabs UK. Oasis HLB cartridges were obtained from Waters.

### Sample preparation

Two standard purified glycopeptides were each resuspended in 50% acetonitrile in water with 0.1% formic acid for direct infusion experiments. To prepare a complex mixture of glycopeptides enriched from a tryptic digest of human serum, we adopted the protocol suggested by Takakura et al.^34^ Briefly, 1 µL of human serum was dissolved in 100 mM ammonium bicarbonate containing 8M urea, reduced by 10 mM TCEP and alkylated by 50 mM 2-CAA, diluted to 6M urea, incubated with 2 µg of LysC for four hours at 37°C, further diluted to 1M urea, and incubated with 2 µg of trypsin at 37°C overnight. After incubation, five-fold volume of ice-cold acetone was added to the peptide mixture and left at -20°C overnight. The resulting mixture was centrifuged at 12000 g, and the supernatant was carefully aspirated and discarded. The pellet containing glycopeptides was redissolved in 5% formic acid in water, desalted using Oasis HLB cartridges and resuspended in 5% formic acid 5% DMSO prior to LCMS analysis.

### Direct infusion and LCMS analyses

All experiments were performed on a Thermo Fisher Scientific Exploris Orbitrap 480 mass spectrometer modified with an Omnitrap platform. The design of direct infusion experiments was described in the previous publication.^32^ Briefly, precursor ions were isolated in the quadrupole mass filter using an isolation mass window of 2 Th, processed in the Omnitrap, and fragments together with unfragmented precursor ions were characterized in the Orbitrap analyzer with a mass resolution of 60,000. The injection times were fixed and set to match the AGC target of 200,000. An ArF ExciStar 200 laser (Coherent, Santa Clara, CA) was used as the source of 193 nm UV light. A FireStar ti60 (Synrad, Mukilteo, WA) laser was used as the source of 10.6 μm IR light with a maximum power output of 60 W. In LCMS analysis of complex mixtures, samples were subjected to LC-MS/MS using an UltiMate 3000 nanoUHPLC system (Thermo Fisher Scientific) coupled to an Exploris-Omnitrap hybrid instrument. The peptides were trapped on a C18 PepMap100 pre-column (300 μm i.d. x 5 mm, 100 Å, Thermo Fisher Scientific) using solvent A (0.1% formic acid in water), then separated on an in-house packed analytical column (50 μm i.d. x 50 cm in-house packed with ReproSil Gold 120 C18, 1.9 μm, Dr. Maisch GmbH) with a gradient of 5% to 45% B (0.1% formic acid in acetonitrile) over 60 min at a flow rate of 100 nL/min. Full scan MS1 survey spectra were acquired in the Orbitrap (scan range 400-2000 m/z, resolution 60,000, AGC target 1,200,000). The 20 most intense peaks identified in a survey scan were selected for fragmentation in the Omnitrap with an AGC target of 200,000 and a maximum injection time of 60 ms, and fragments together with unfragmented precursor ions were characterized in the Orbitrap analyzer in a single scan with a mass resolution of 45,000. SceHCD experiments were performed in the HCD cell of Exploris at 20, 30, and 40% of normalised collision energy (NCE) using an AGC target of 100,000, a maximum injection time of 60 ms, and a mass resolution of 30,000. The first mass in MS2 scans was locked at 125 *m/z* in all experiments. The low-mass cutoff of the Omnitrap was held constant at 150 *m/z* during acquisition.

### Data analysis

Raw LCMS data were analysed in the FragPipe 23.1 platform using the MSFragger 4.3 search engine^35^ featuring a glycopeptide search module.^36^ The search was performed using “glyco-N-HCD” or “glyco-N-Hybrid” default parameters with, depending on the purpose of the search, various fragment ion series specified for identification in the “MSFragger” tab. Variable modifications included oxidation of methionine and protein N-terminal acetylation, and cysteine carbamidomethylation was set as fixed modification. Glycans were searched within the “Human_N-glycan_Medium” database. Results were filtered to 1% FDR on both peptide and glycan levels.^37, 38^

### Spectra annotation and data availability

Selected spectra were manually annotated using fragment *m/z* values generated *in-silico* in Protein Prospector v6.5.0 (http://prospector.ucsf.edu) and GlycoWorkbench v1.2.4105.^39^ Nomenclature for the annotation of spectra used in this paper is given in Supplementary Figure S1. The raw files were deposited into the PRIDE repository with the project identifier PXD071023.

## Results and discussion

### Initial characterisation of EID, UVPD for glycopeptides

The distinct fragmentation behaviour of glycopeptides requires investigation of what are its optimal parameters, more specifically, the number and energy of laser pulses for UVPD and irradiation time for EID. We started with the study of standard glycopeptides KVANKT modified with glycoforms HexNAc4Hex5NeuAc2 or HexNAc4Hex3Fuc (Supplementary Figure S1d) in direct-infusion UVPD and EID experiments. In three series of experiments, we subjected the glycopeptides to i) UV radiation using various numbers of pulses at fixed pulse energy, ii) various pulse energies using the fixed number of pulses, and iii) to high-energy electrons with a range of irradiation times. We found that both UVPD and EID generate an abundance of fragments of different types, including peptide, glycosidic and cross-ring fragments. When comparing the glycoforms we observed that the glycan composition affects the sequencing of the peptide: the presence of additional galactose and sialic acid saccharides on both antennas reduces the number of peptide backbone fragments, in particular on the C-terminus irrespective of applied conditions (Supplementary Figures S2, S3). Interestingly, it is galactose and sialic acid that demonstrated extensive cross-ring dissociation (A- and X-types of glycan fragments) in both EID and UVPD, while the HexNAc4Hex3Fuc glycoform shows almost exclusively ^1,5^X fragments of various saccharides which is simply the combination of an anomeric carbon and one oxygen atom. While UVPD and EID indeed produce a plethora of cross-ring and glycosidic internal glycan fragments, many of them (labelled with asterisk in Supplementary Figures S2-S5) have identical masses which makes it impossible to determine their identity using standard analytical workflows. In addition, we were able to discover some peaks which are present in the spectra of both glycoforms of the standard glycopeptides but can only be annotated for one of them. For example, Supplementary Figure S6 shows a zoomed-in EID and UVPD spectra of standard glycopeptides around the m/z 382.134. While it is tempting to assign this peak to the [^0,3^A_Gal_+NeuAc] fragment in the spectrum of the HexNAc4Hex5NeuAc2 glycopeptide, the question remains about its identity in the spectrum of the HexNAc4Hex3Fuc glycoform. The solution we propose is to consider secondary cross-ring fragmentation. This way, the peak can be assigned to [^0,3^A_Man_+Man+^3,5^X_GlcNAc_] or [^1,4^A_Man_+Man+^3,5^X_GlcNAc_] (Supplementary Figure S6). We acknowledge however, that such a solution will further complicate an already complex analysis of a glycopeptide spectra.

The charge state has hardly any effect on the fragment generation (Supplementary Figures S4, S5), possibly suggesting a charge-remote mechanism of dissociation. In terms of fragment diversity, EID produces noticeably more cross-ring fragments, while UVPD performs slightly better at dissociating the peptide backbone. Importantly, both techniques cleaved all glycosidic bonds producing complete sequences of Y fragments for both glycoforms and retained the core fucose in the HexNAc4Hex3Fuc. To determine the effect of the experimental parameters on the fragmentation efficiency, we plotted the absolute intensities of representative peptide, glycosidic, and cross-ring fragments against the number and energy of laser pulses in UVPD (Supplementary Figures S7, S8) and irradiation length in EID (Supplementary Figure S9). We observed a continuing increase of fragmentation intensity within the range of chosen values (Supplementary Figures S7-S9). It is noteworthy that the non-modified peptides experienced optimal fragmentation intensity well within this range of values.^33^ This suggests that higher energy density may be required to dissociate glycopeptides compared to non-modified peptides but further investigation with a larger pool of glycopeptides is necessary. Reassuringly, our direct-infusion EID of the HexNAc4Hex5NeuAc2 glycopeptide and those by Wei and co-workers^18^ produced similar spectra suggesting EID on an Omnitrap can be reproducible and is potentially independent of the mass analyser.

### LCMS experiments for all fragmentation techniques

In order to characterize ECD, AI-ECD, UVPD and EID on a larger range of glycopeptides, we extended our study to LCMS analysis of complex mixtures. We generated a tryptic digest of human serum that had its glycopeptide population enriched by acetone precipitation.^34^ Such treatment preferentially removes hydrophobic compounds yielding a mixture of mainly hydrophilic peptides and N-glycosylated peptides. The choice of the search engine for LCMS data analysis was dictated, first, by the need to control false discovery rate (FDR) on both peptide and glycan levels, and second, by the need to be able to freely select the types of peptide fragment ions for analysis of EID and UVPD data. We therefore opted for the MSFragger search engine^35^ implemented as a part of the FragPipe proteomics platform. FragPipe provides a tool for glycopeptide analysis *via* its MSFragger-Glyco module,^36^ enables glycan FDR control,^38^ and allows users to customize peptide fragment ion types used for identification.

We first performed a stepped high-energy collisional dissociation (sceHCD) analysis of the glycopeptide mixture at 20, 30 and 40% normalized collisional energy and searched for N-glycopeptides. This method reflects a common approach to study N-glycopeptides and will serve as a benchmark for the other types of glycopeptide dissociation. In this experiment, we identified 4314 non-glycopeptide spectrum matches (non-glycoPSMs), 2104 glycoPSMs, 2007 unique non-glycosylated peptides, and 1256 unique glycopeptides which equates to 33% enrichment on the (glyco)PSM level and 38% — on the unique (glyco)peptide level. In line with earlier studies,^40, 41^ the distribution of precursors *m/z-*binned values shows an apparent separation between the two groups where glycopeptide precursors are shifted towards higher *m/z* (Supplementary Figure S10).

We then investigated the efficiency of ECD in the analysis of the glycopeptide mixture. In total, 2025 PSMs were identified in a 100 ms (irradiation time) ECD experiment with only 131 of them being glycoPSMs. Supplementary Figure S11 shows a spectrum of [DGTGHGNSTHHGPEYMR]^5+^ peptide modified with [HexNAc4Hex5NeuAc2]. The spectrum contains *c* and *z* fragments resulting in a complete sequence coverage of the peptide. However, the extent of fragmentation of glycosidic bonds is modest since low-energy electrons dissociate primarily C_α_-N bonds within the amino-acid backbone and add very little vibrational energy.^42^ As most search engines for glycopeptide LCMS analysis rely on the presence of oxonium and *B/Y* ions in a spectrum, ECD without supplemental activation consequently results in relatively small numbers of glycoPSMs. Coon and co-workers have shown that supplemental co-activation by IR radiation can be beneficial for ETD analysis of glycopeptides (AI-ETD).^24^ We reasoned that, similar to that study, the combination of ECD and IRMPD within one MS scan (activated-ion ECD, AI-ECD^43, 44^) may lead to the increase in the number of identified glycoPSMs by providing glycan fragments and at the same time improving the amino acid sequence coverage by *c* and *z* fragments *via* vibrational activation of charge-reduced products. To achieve the best results, the energy supplied by the IR laser is required to be sufficiently low to minimise secondary fragmentation of *c,z* ions generated by ECD. To facilitate efficient AI-ECD, we therefore moved the IR laser focus out of the ECD reaction chamber by adjusting the optical lens and limited the laser co-irradiation time to 50 ms at 13% laser duty cycle while keeping the duration of the electron irradiation at 100 ms. The effect of ion activation is demonstrated in Figure 1 showing ECD and AI-ECD spectra of the peptide [QQQHLFGSNVTDCSGNFCLFR]^3+^ modified with a complex glycan [HexNAc4Hex5NeuAc2]. ECD results in only five peptide and two glycan fragments of the precursor (Figure 1a) while AI-ECD yields a near complete peptide sequence coverage and dissociates four glycosidic bonds within the glycan (Figure 1b). Interestingly, the AI-ECD spectrum contains many *b,y* peptide fragments which is not typical for the low energy IRMPD of glycans.^10, 11, 20^ In total, we identified 473 glycoPSMs in AI-ECD experiments, a significant improvement over ECD only. Studying distribution of precursors (Figure 1c) showed that the additional vibrational energy only slightly improved the identification rate at smaller *m/z* bins but greatly enhanced the numbers at higher *m/z*. The IR-activation appears to be most productive for 3+ and 4+ glycopeptide precursors, as very few doubly charged precursors were identified in both ECD and AI-ECD (Figure 1d-g). An extensive loss of an acetyl group from charge-reduced species, competing with the formation of informative fragments such as *c,z* ions, can be observed in all ECD mass spectra demonstrated in this study (Figure 1a,b and Supplementary Figure S11). An attempt was made previously to describe the loss of an acetyl group in terms similar to ECD of non-modified peptides.^8^ The proposed mechanism involves an attack of the N-acetyl nitrogen atom in a glycan by a hydrogen atom liberated in ECD and a formation of an N-centred hypervalent radical which subsequently fragments by loss of an acetyl radical.^8^

**Figure 1.**
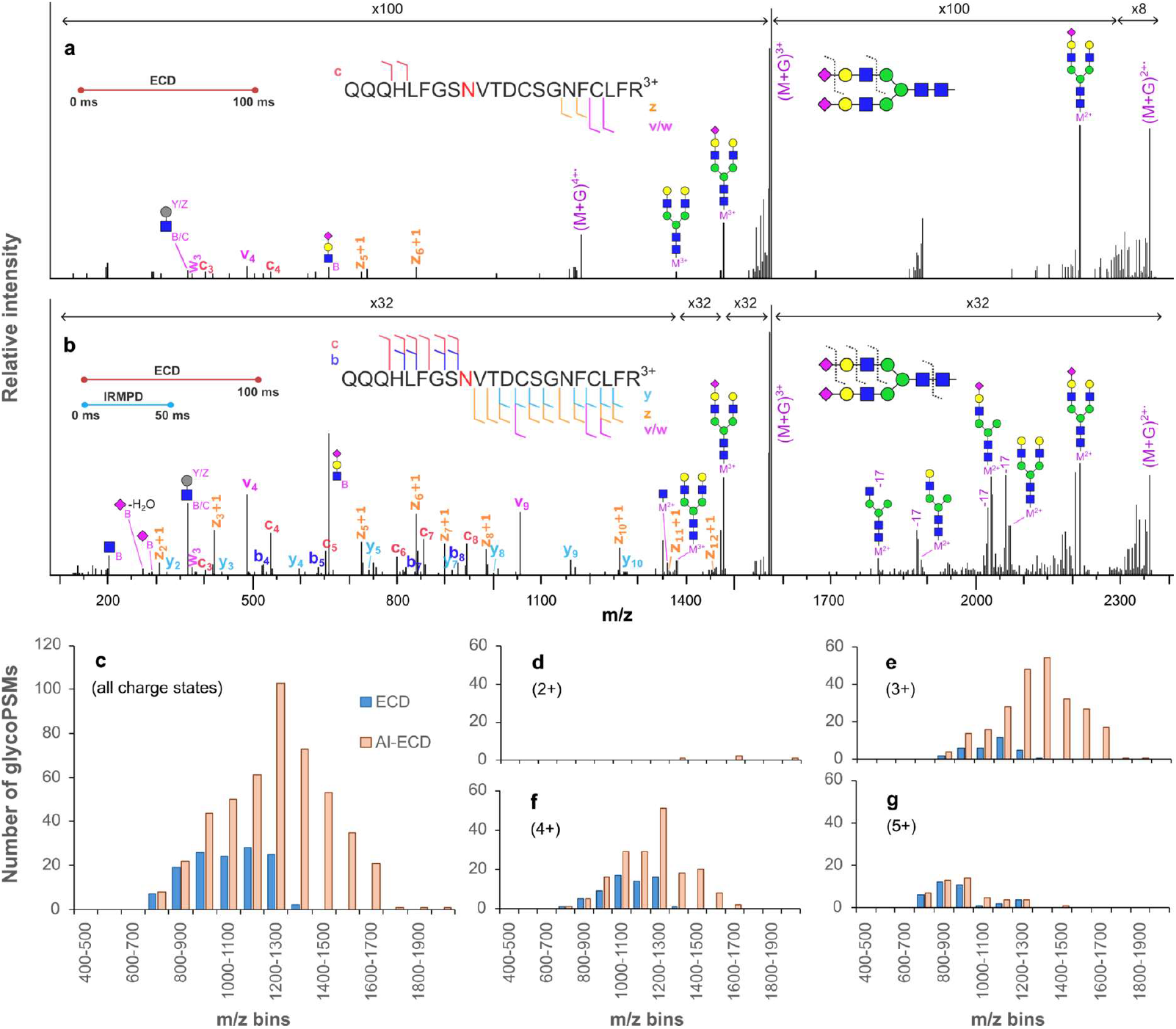
ECD **(a)** and AI-ECD **(b)** spectra of N-glycosylated QQQHLFGS<u>N</u>VTDCSGNFCLFR^3+^ acquired in LCMS of a complex glycopeptide mixture. Intact peptide is denoted as M, and precursor ions are annotated as (M+G). All annotations of intact peptides retaining partial glycan correspond to *Y* fragments unless otherwise specified. Precursor ions were irradiated by electrons for 100 ms. In AI-ECD, precursor ions were in addition co-irradiated by IR light at 13 % of the laser duty cycle for the first 50 ms of ECD. **c-g:** M/z-binned distributions of the numbers of glycoPSMs identified in ECD (blue) and AI-ECD (orange) analyses of complex glycopeptide mixture: (**c)** all charge states, **(d)** doubly, **(e)** triply, **(f)** quadruply, **(g)** 5+ charged precursors; *b,y,c,z* fragments were used for identification.

Next, we optimized the parameters of EID and UVPD in LCMS experiments. We recorded the numbers of non-glycoPSMs and glycoPSMs while varying the number or energy of laser pulses in UVPD and irradiation time in EID and keeping other parameters fixed. For automated data analysis, we compared the following combinations of peptide fragments: *b,y*, as they were found to be essential for analysis of UVPD and EID of non-modified peptides,^33^ *a,b,y*, as *a* ions are the third most frequent type of ions in UVPD and EID,^33^ *b,y,c,z* as default parameters in the “N-Hybrid” workflow in MSFragger, and we also included *a,b,y,c,z* set of fragments specifically in the analysis of EID data, as this combination showed the best results for non-modified peptides.^33^

The analysis of the glycopeptide EID and UVPD datasets showed the presence of both non-modified and N-glycosylated peptides (Figure 2a,b). The identification rates of non-glycoPSMs at different UVPD and EID parameters are in line with our previous study.^33^ The identification of glycoPSMs, however, follows completely different patterns, especially in UVPD. The numbers of identified glycoPSMs across the range of UVPD and EID parameters depends on which peptide fragments are used for analysis (Figure 2b). The *b,y* pair of fragments showed clearer trends in their results in UVPD with only one maximum and no sharp increases/drops in the range of energies and numbers of laser pulses, while simply adding *a* ions to the search parameters causes the appearance of one minimum and two maxima (Figure 2b). Furthermore, the optimal fragmentation in the LCMS UVPD analysis of glycopeptides is achieved at remarkably higher energies and pulses than those required to identify non-modified peptides. In the analysis using *b,y* types of peptide fragments, two laser pulses at 6 mJ/pulse allowed to identify only 4 glycoPSMs, and this number gradually increases until the maximum of 555 glycoPSMs is reached at 8 laser pulses (Figure 2b). For non-glycoPSMs this dependency flattens out after 4 laser pulses (Figure 2a), in agreement with our findings in the UVPD-LCMS of non-modified peptides.^33^ In a range of pulse energies, the number of identified glycoPSMs starts with 365 at 4 mJ/pulse and reaches the maximum of 799 at 8 mJ/pulse. These observations are in line with the findings described for the UVPD of glycopeptides in the group of Brodbelt.^23, 25^ As the search engine used in this work employs “peptide first” approach to identify glycopeptides, the identification of a precursor relies on the availability of peptide fragments in the first place. We assume therefore that the efficiency of the fragmentation of the peptide chain in a glycopeptide is significantly lower than in a non-modified peptide. To explain this phenomenon in UVPD, we note that a 193 nm photon can be absorbed either by an amide bond within the peptide backbone^45, 46^ or by N-acetylhexosamine saccharides.^47^ The absorption of a photon by the latter may lead to an intensive loss of an acetyl moiety (Supplementary Figures S2, S4), cleavage of labile glycan bonds due to intramolecular vibrational redistribution, or cross-ring fragmentation via unknown mechanisms. Similar reasonings can be applied to the EID of glycopeptides, even though its initiation mechanism is fundamentally different. Depending on the location and kinetics of the interaction between an electron and a glycopeptide ion, the ion may undergo dissociation at either peptide backbone, amino acid sidechain, or anywhere within the glycan. We conclude therefore that glycans, especially those that are ‘big and complex’, shield the backbone from electrons or photons which leads to lower fragmentation rates of the peptide backbone.

**Figure 2.**
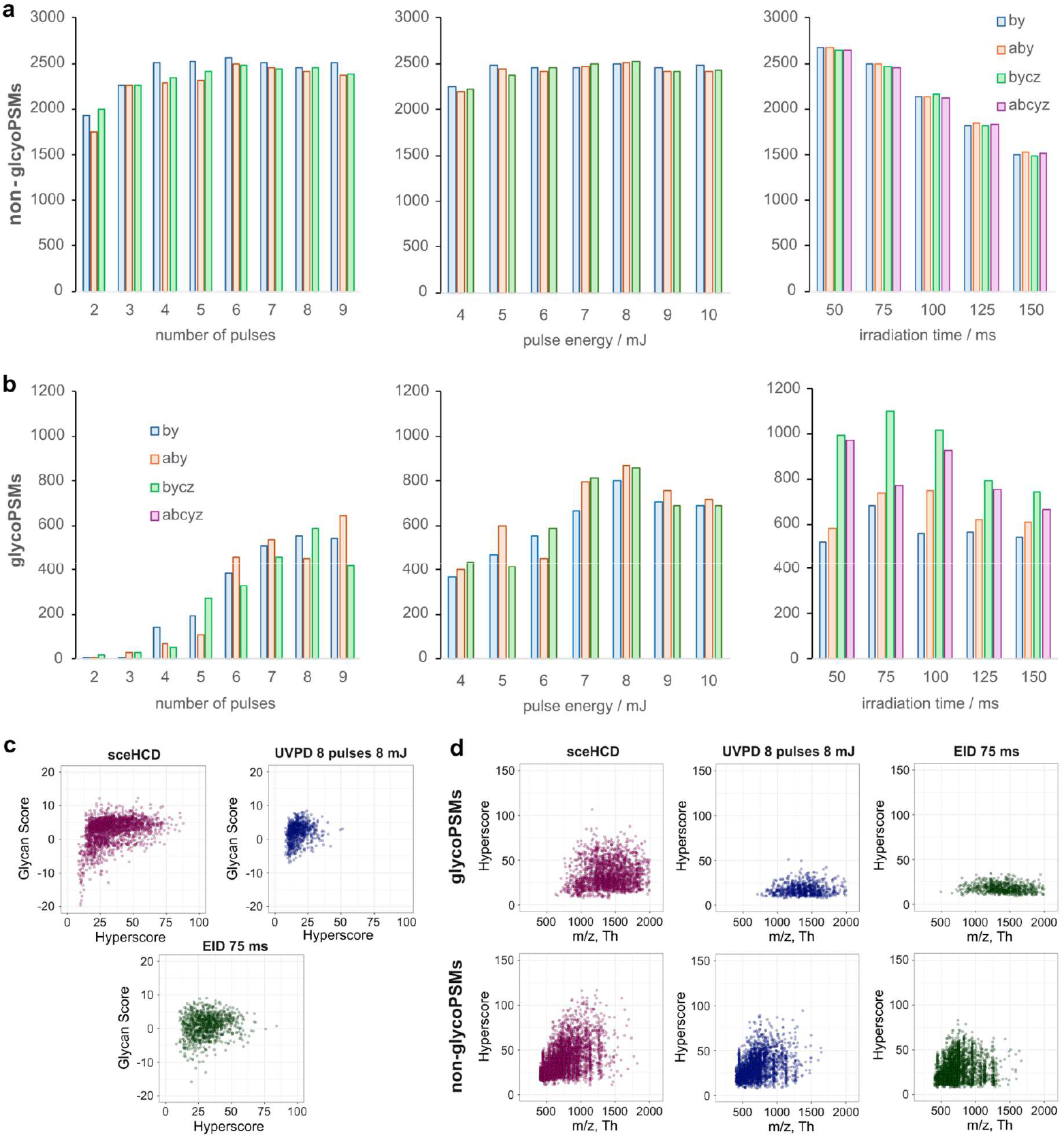
Numbers of identified non-glycoPSMs (**a**) and glycoPSMs (**b**) in UVPD in a range of numbers of laser pulses at 6 mJ/pulse (left) and pulse energies for 8 pulses (middle) and in EID data in a range of electron irradiation times (right). **c**) Glycan scores plotted against hyperscores for glycoPSMs identified in sceHCD experiments, UVPD experiments using *b,y* peptide fragments for analysis and in EID experiments using *b,y,c,z* peptide fragments for analysis. **d**) Hyperscores plotted against *m/z* values of glycoPSMs (top) and non-glycoPSMs (bottom) identified in sceHCD, UVPD and EID experiments using *b,y* peptide fragments for analysis.

In EID, *b,y,c,z* showed noticeably better results in terms of identification numbers than *b,y* and *a,b,y* combinations (Figure 2b). This is a somewhat unexpected observation, as no such difference between different combinations of peptide fragments was observed for non-glycoPSMs in the EID data and in the UVPD data on both glyco- and non-glycoPSM levels (Figure 2a,b). This implies that EID of glycopeptides produces on average more *c,z* types of peptide fragments than UVPD. The number of non-glycoPSMs drops as the irradiation time increases, potentially because of lower scan rates (Figure 2a). When using *b,y,c,z* peptide fragments for analysis, the number of glycoPSMs reaches its maximum of 989 at 75 ms and then begins to decline also likely due to lower scan rate (Figure 2b).

To assess how both techniques perform in sequencing of peptide and glycan parts of glycopeptides, we plotted the distributions of two types of scores generated by MSFragger – Glycan score and hyperscore – against each other for glycoPSMs acquired at optimal parameters of sceHCD, UVPD and EID (Figure 2c, Supplementary Figure S12). We also plotted the distributions of hyperscores for glyco- and non-glycoPSMs against precursor *m/z* identified in the same experiments (Figure 2d, Supplementary Figure S13). Glycan score reflects the abundance of Y-type glycan fragments and oxonium ions,^38^ and hyperscore correlates with the number of peptide fragments.^35^ In sceHCD, the distribution of Glycan scores is by design centred at zero,^48^ with higher (positive) values indicating higher number of matched glycan-related peaks.^35^ Our sceHCD data was heavily biased towards higher Glycan Scores and has a long tail of IDs with low values of both scores (Figure 2c). We were pleased to see that UVPD and EID in our experiments produced very similar distributions with regards to Glycan score (Figure 2c, Supplementary Figure S12), which suggests sufficient fragmentation of glycosidic bonds to form abundant oxonium and Y-ions in these two processes. The hyperscores, on the other hand, are drastically different in sceHCD, where they reach the values of around 75, and in UVPD and EID, where they largely fall below 25 (Figure 2d, Supplementary Figure S13a) when *b,y* peptide fragments were used for data analysis. Collisional dissociation is known to produce partially and fully deglycosylated peptide fragments^49,50^ that contribute to the hyperscore in sceHCD. Relatively low hyperscores in UVPD and EID suggest that these two processes generate fewer identifiable peptide fragments, in particular C-terminal from the glycosylation site for *x,y,z* types of fragments, and N-terminal — for *a,b,c* types of fragments. Finally, the distribution of hyperscores of glycopeptides in EID spans higher values than in UVPD when *b,y,c,z* ions are used for analysis, supporting our earlier statement that EID produces more *c,z* types of peptide fragments (Supplementary Figure 13b,c).

After we performed this first round of optimization, we compared the quality of UVPD, EID, and sceHCD LCMS spectra of a typical glycopeptide. To this end, we annotated respective spectra of a triply charged peptide AALAAFNAQNNGSNFQLEEISR modified with a complex N-glycan [HexNAc4Hex5NeuAc2]. These spectra were acquired in the experiments where optimal parameters of fragmentation were used, namely, 8 pulses at 8 mJ/pulse for UVPD and 75 ms of irradiation time for EID. Noteworthy, the amount of information we observed in this UVPD and EID LCMS spectra (Figure 3) is on par with direct-infusion experiments of glycopeptide standards (Supplementary Figures S2-S5) which suggests that this instrument design is suitable for detailed structural characterization of glycopeptides on the LCMS scale. As expected, stepped collisional fragmentation readily generates a complete peptide sequence coverage by (deglycosylated) *b,y* ions and dissociates all glycosidic bonds within the glycan. UVPD and EID also confirm the peptide identification by providing an abundance of main-series peptide fragments. We didn’t find any peptide fragments containing an intact glycan, and we only observed one deglycosylated *b*-ion in UVPD, which explains lower hyperscores observed in EID and UVPD of glycopeptides compared with sceHCD. All three spectra contain cross-ring fragments (Figure 3), which are useful in that they may allow one to determine types of glycosidic bonds.

**Figure 3.**
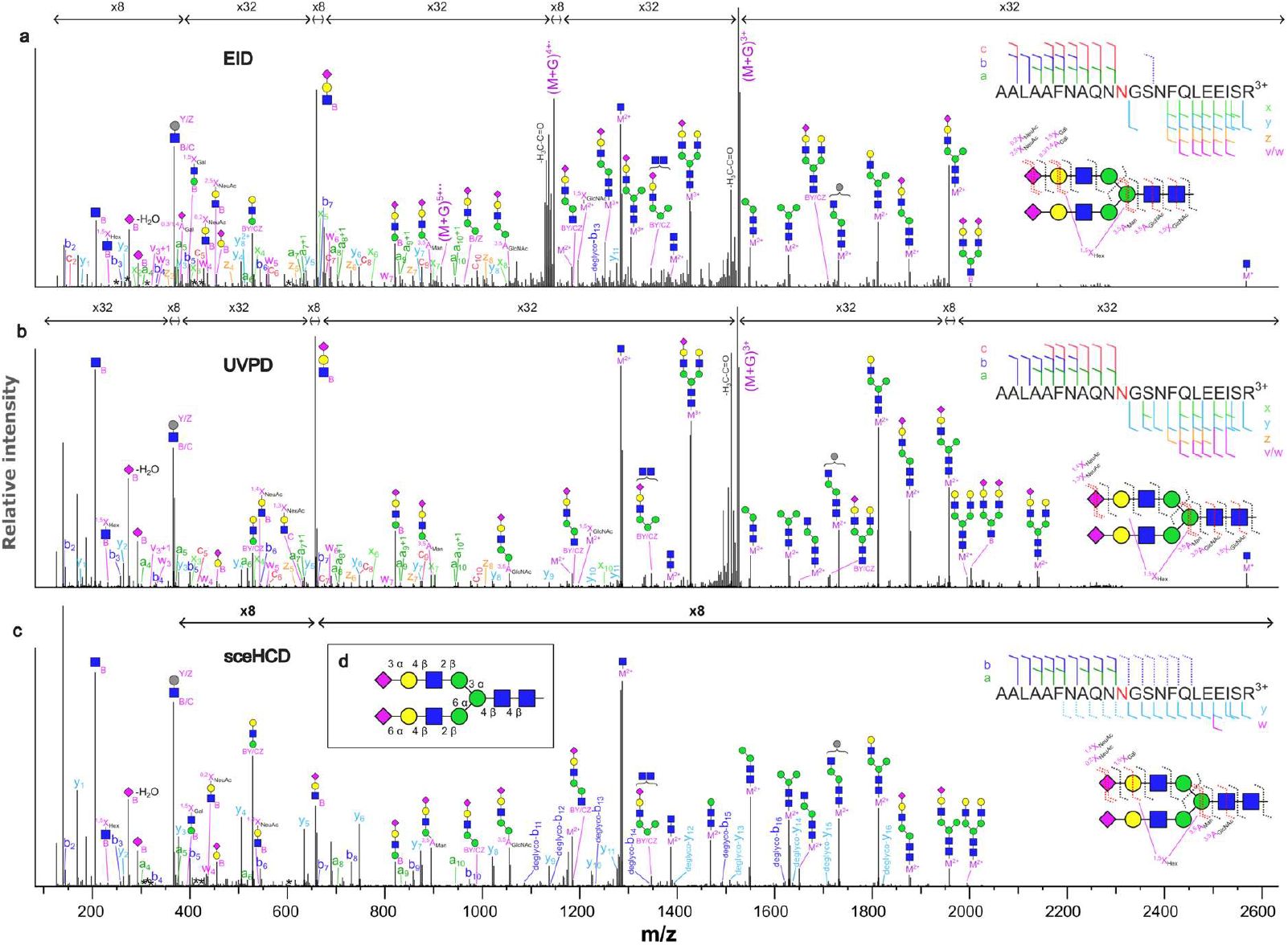
UVPD, EID, and sceHCD LCMS spectra of an N-glycopeptide. EID **(a)**, UVPD **(b)**, and sceHCD **(c)** spectra of N-glycosylated AALAAFNAQNNGSNFQLEEISR^3+^ peptide acquired in the LCMS analysis of a complex glycopeptide mixture. M/z values matching to multiple isobaric internal glycan fragmentation products are marked with asterisks. Intact peptide is denoted as M, and precursor ions are labelled as (M+G). All annotations of intact peptides retaining partial glycan correspond to *Y* fragments unless otherwise specified. Dashed labels correspond to fully deglycosylated peptide fragments. For UVPD, 8 pulses at 8 mJ/pulse were used; in EID, precursor ions were irradiated by 25 eV electrons for 75 ms. Assumed composition of the glycan is shown in (**d**).

As discussed above, the increase in energy required to fragment a glycopeptide over a non-modified peptide suggests that the size of the glycan may influence the extent of fragmentation. We hypothesized that a peptide modified with a relatively small glycan has a higher chance to be retained upon EID or UVPD than a larger glycan. Figure 4 shows EID, UVPD, and sceHCD LCMS spectra of a triply charged 21-amino-acid long O-glycopeptide TEHLASSSEDSTTPSAQTQEK with nine potential sites of O-glycosylation. SceHCD provides an excellent peptide and glycan sequence coverages by *b, y*, and *Y* fragments; it, however, exhibits substantial loss of the sugar leaving an intact amino acid behind which precludes site localisation. The only two peptide fragments retaining a part of the glycan are b_13_ and b_14_ ions bearing a GalNAc fragment, which narrows down the list of possible glycosylation sites to seven amino acids (Figure 4c). UVPD and EID spectra contain a rich array of peptide main series fragments in addition to few products of glycosidic bond and cross ring dissociation. Importantly, y_9_^2+^, b_13_^2+^ and a range of *a*-type fragments allow to localize the O-glycan to Thr13 residue (Figure 4). In addition, the presence of the fragment d_13_^2+^ (in both EID and UVPD) might possibly serve as indirect evidence of the presence of a modification at Thr13.

**Figure 4.**
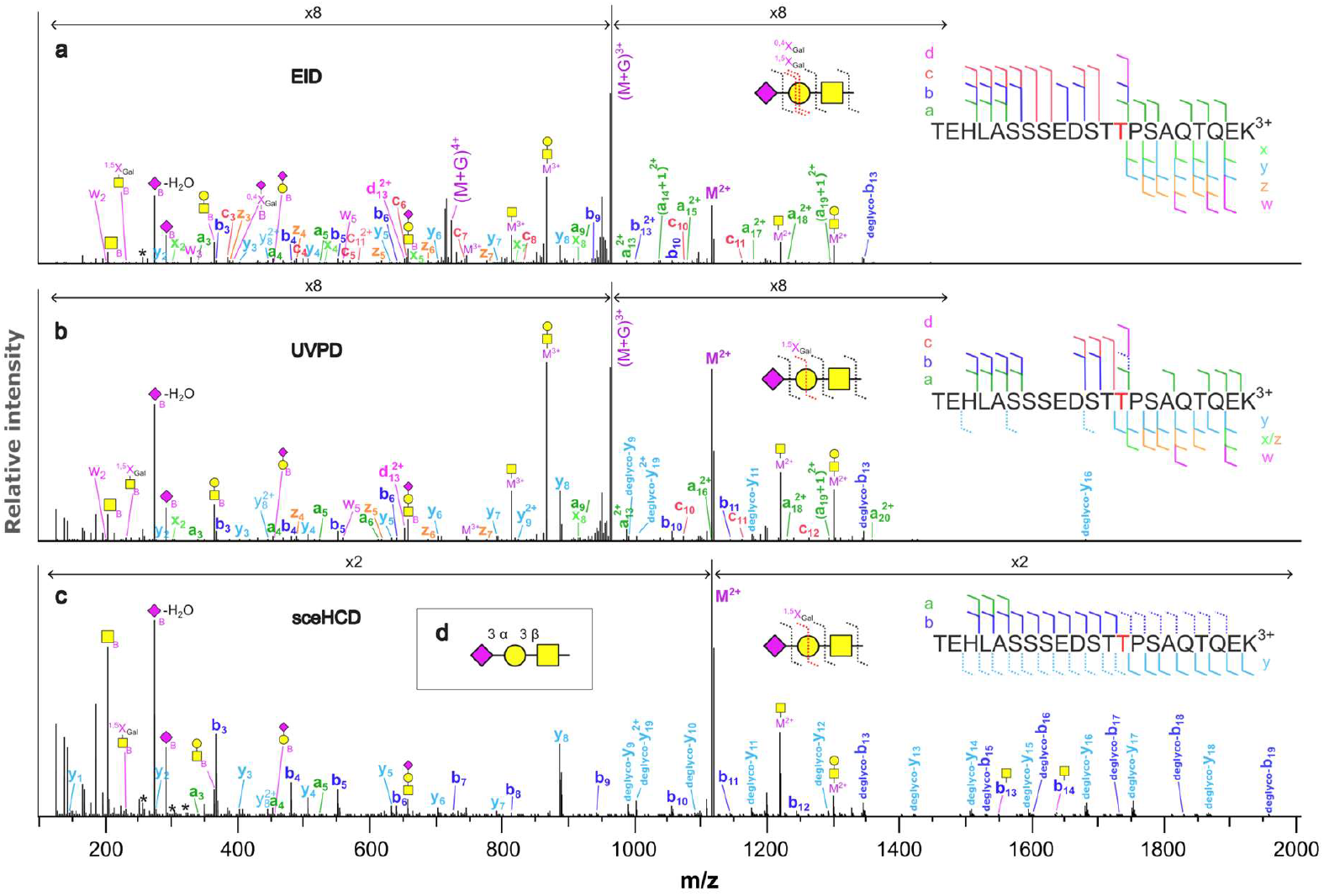
UVPD, EID, and sceHCD LCMS spectra of an O-glycopeptide. UVPD **(a)**, EID **(b)**, and sceHCD **(c)** spectra of O-glycosylated TEHLASSSEDST<u>T</u>PSAQTQEK^3+^ acquired in the LCMS analysis of a complex glycopeptide mixture. M/z values matching to multiple isobaric internal glycan fragmentation products are marked with asterisks. Intact peptide is denoted as M, and precursor ions are annotated as (M+G). All annotations of intact peptides retaining partial glycan correspond to *Y* fragments unless otherwise specified. Dashed labels correspond to fully deglycosylated peptide fragments. For UVPD, 8 pulses at 8 mJ/pulse were used; in EID, precursor ions were irradiated by 25 eV electrons for 100 ms. Assumed composition of the glycan is shown in (**d**).

We also compared the absolute numbers of identifications using optimal fragmentation parameters and MS1 *m/z* range of 900-2000 Th to minimize interference with non-glycosylated peptides. Table 1 shows the numbers for all fragmentation techniques studied here. SceHCD predictably produced the highest number of identifications, nearly twice as many as the second best, largely due to high scan rate and the ability to adjust collision energy for each precursor *m/z* and charge state. UVPD has the same efficiency per MS2 scan as sceHCD, 6.8% vs 6.7%, and EID falls behind with only 5.9%.

**Table 1.** Maximum numbers of non-glycoPSMs and glycoPSMs identified in a complex glycopeptide mixture by each fragmentation technique.

|  | fragmentation parameters | non-glycoPSMs | glycoPSMs | number of MS2 scans | glycoPSMs per MS2 scan, % | peptide fragment types used for data analysis |
| --- | --- | --- | --- | --- | --- | --- |
| sceHCD | 20, 30, 40% NCE | 1480 | 2749 | 40805 | 6.7 | b,y |
| ECD | 100 ms irradiation | 305 | 206 | 14683 | 1.4 | b,y,c,z |
| AI-ECD | 50 ms co-irradiation, then 50 ms just ECD | 474 | 1045 | 14834 | 7.0 | b,y,c,z |
| UVPD | 8 pulses, 8 mJ/pulse | 893 | 1383 | 20270 | 6.8 | b,y |
| EID | 75 ms irradiation, 25 eV | 482 | 948 | 16100 | 5.9 | b,y,c,z |

ECD alone is not a very efficient method for glycopeptide analysis in that it is not able to produce sufficient amount of oxonium and Y-type ions needed for identification. It, however, has arguably higher potential than EID and UVPD for sequencing peptide backbones while retaining intact glycans (compare, for example, Figure 3 and Supplementary Figure S11). Supplemental IR activation boosts the numbers of glycoPSMs identified in ECD experiments more than five times which is enough to overcome EID but not UVPD (Table 1). It is worth mentioning though that the efficiency of both UVPD and EID is much less charge-dependent than ECD which implies adjusting electron irradiation time to the charge state of each precursor might be required for the latter. In addition, the efficiency of IRMPD depends too on the charge state and *m/z* (which are proxy for size) of a glycopeptide precursor.^20^ While insufficient activation results in too few oxonium and glycosidic fragments, too much IRMPD may lead to adverse fragmentation of *c,z* fragments generated by ECD. Both these outcomes are undesirable, and finding a sweet spot to enable data-dependent approach can be challenging but highly beneficial for ECD and AI-ECD of glycopeptides. Furthermore, as the scan rate in sceHCD is considerably higher than in any other technique employed here (Table 1), we believe that the identification numbers can be further increased by implementing product-dependent approach, where ExD/UVPD event can be triggered only following a detection of some oxonium ions in a “scout” HCD scan.

## Conclusion

This study establishes the Omnitrap as a potent technology that allows for acquisition of high-quality detailed UVPD and EID spectra of complex N-glycopeptides in both direct-infusion and LCMS modes. Through systematic evaluation of UVPD and EID parameters, we show that alternative high-energy activation modes provide fragmentation richness and structural information that extend beyond what collisional methods can achieve, but at the same time highlight the need for development of optimised acquisition strategies for optimal sequencing of both peptide and glycan parts of the analyte. We furthermore show that ECD alone suffers from low dissociation yields which can be rectified by supplemental activation by IR light. Finally, we demonstrate that the scoring patterns produced by a search engine for EID and UVPD glycopeptide data are substantially different compared to collisional data which suggests further software development may be required for efficient analysis of such types of data.

## Acknowledgements

The Next Generation Chemistry group at the Franklin Institute is supported by the EPSRC (V011359/1 (P)). The authors express their gratitude to Alexey Nesvizhskii and Daniel Polasky from University of Michigan for sharing insights into the functionality of MSFragger search engine, to the members of Fasmatech and MS Vision teams for their continuous technical support as well as to Ajay Jha, Ben Davis and Weston Struwe from University of Oxford for their critical comments on the manuscript draft.

## Author contributions

N.L. collected and analysed the data. N.L. and S.M. conceptualized and designed the experiments, interpreted the data, and wrote the manuscript.

