## Supporting Information for "Evaluating the performance of photon- and electron-based fragmentation methods in Omnitrap-LCMS analysis of N-glycopeptides"

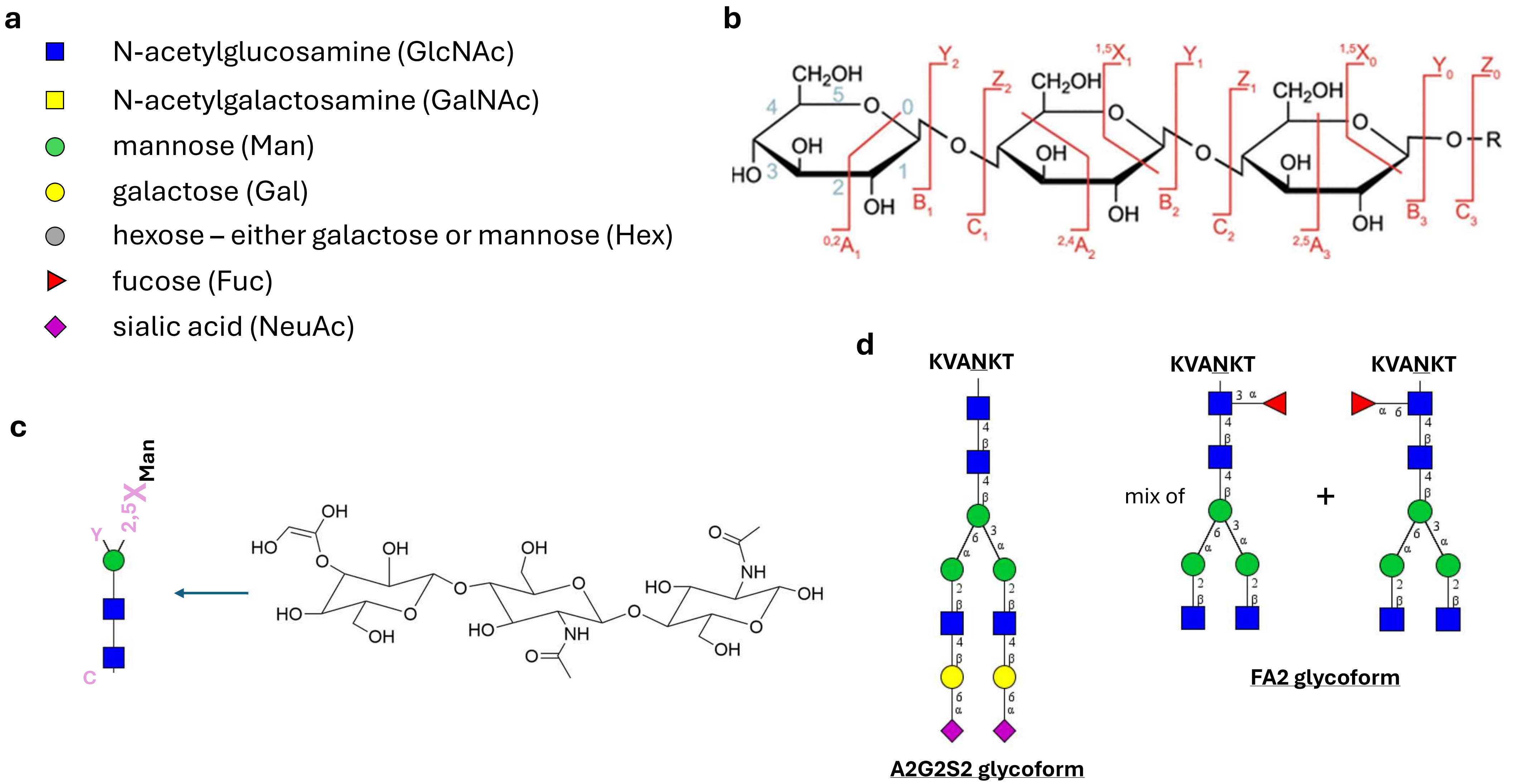


**Supplementary Figure S1**. **a)** Monosaccharide symbol nomenclature, **b)** glycan fragmentation nomenclature as proposed by Domon and Costello [Domon, Costello *Glycoconjugate J.*, 5 (1988), p. 397], figure adopted from [Grabarics, Pagel et al., *Chem. Rev*. 2022, 122, 8, 7840-7908], **c)** example of annotation of an internal glycan fragment used throughout this paper, **d)** compositions of two standard glycopeptides used in this work in direct infusion experiments.

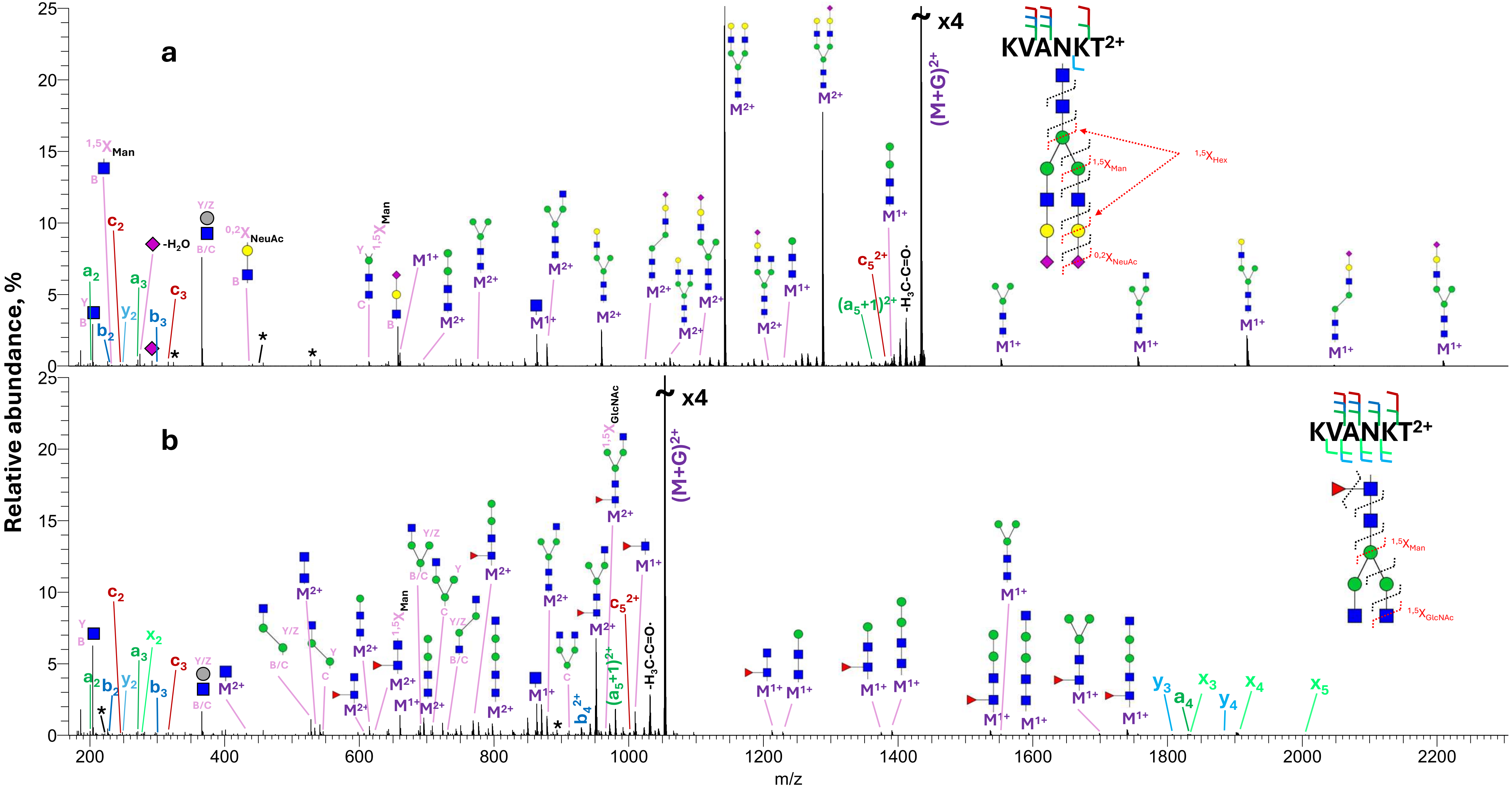


**Supplementary Figure S2**. Averaged MS2 UVPD mass spectra of doubly charged A2G2S2 (**a**) and FA2 (**b**) glycoforms of the standard glycopeptide, acquired using 10 laser pulses and 10 mJ/pulse. M/z values matching to multiple isobaric internal glycan fragmentation products are marked with asterisks. Intact peptide is denoted as M, and precursor ions are annotated as (M+G). All annotations of intact peptides retaining partial glycan correspond to Y fragments. Fragments matching neutral losses of water and –CO were not annotated.

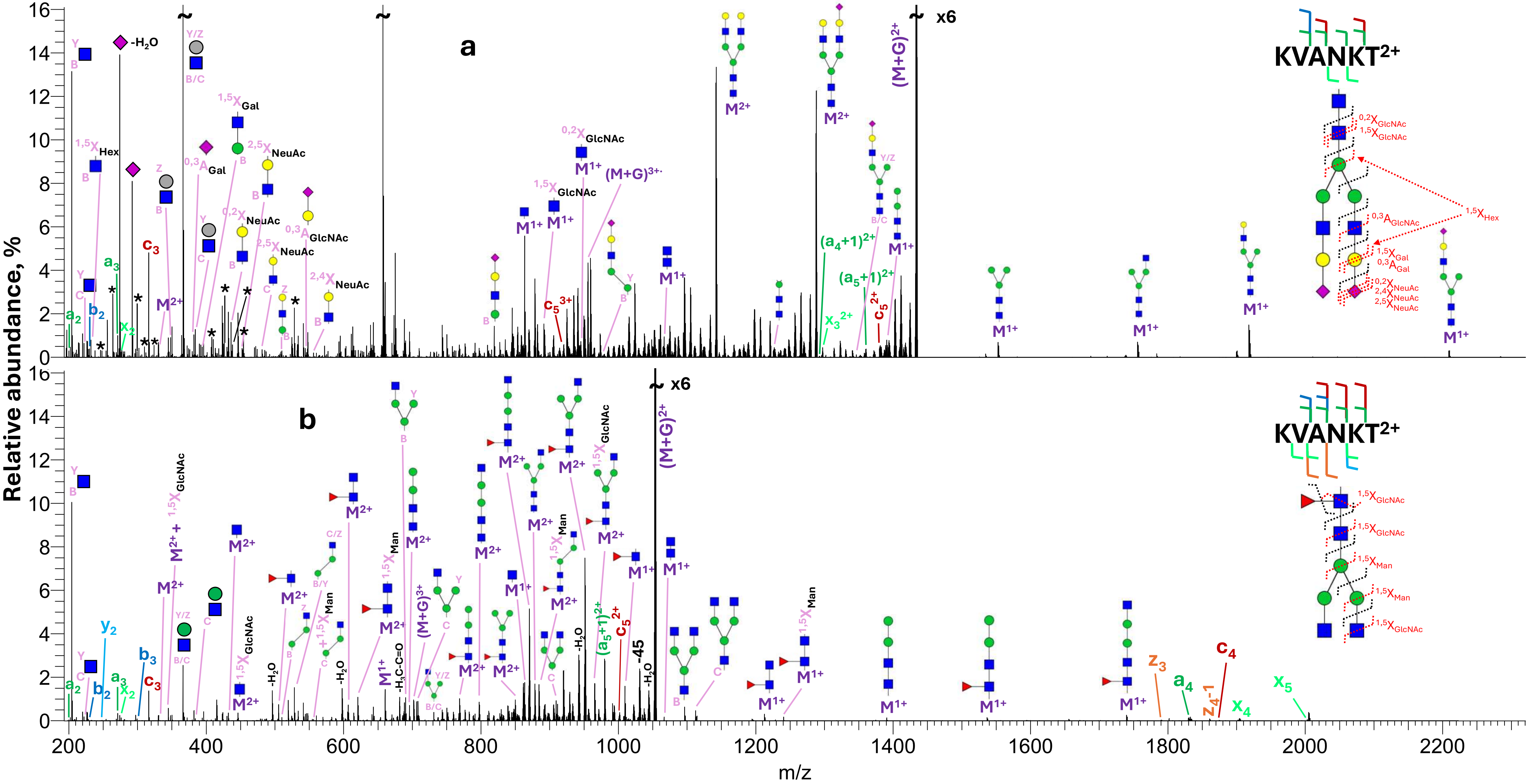


**Supplementary Figure S3**. Averaged MS2 EID mass spectra of doubly charged A2G2S2 (**a**) and FA2 (**b**) glycoforms of the standard glycopeptide, acquired after irradiating the precursor ions for 150 ms by 25 eV electrons. M/z values matching to multiple isobaric internal glycan fragmentation products are marked with asterisks. Intact peptide is denoted as M, and precursor ions are annotated as (M+G). All annotations of intact peptides retaining partial glycan correspond to Y fragments unless otherwise specified. Fragments matching neutral losses of water were not labelled. Typically, only one charge state of a fragment was annotated.

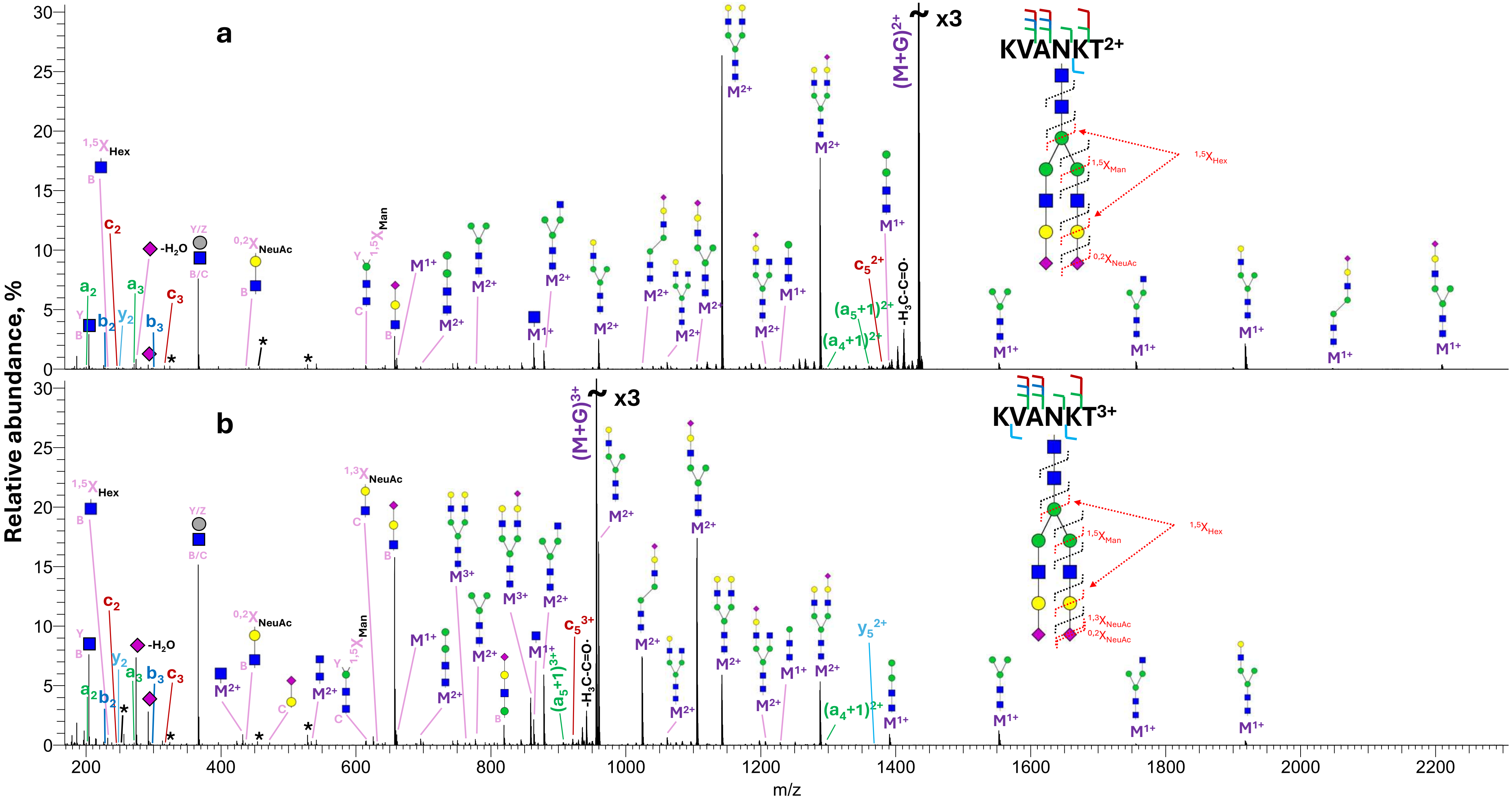


**Supplementary Figure S4**. Averaged MS2 UVPD mass spectra of doubly (**a**) and triply (**b**) charged A2G2S2 glycoform of the standard glycopeptide, acquired using 10 laser pulses and 10 mJ/pulse. M/z values matching to multiple isobaric internal glycan fragmentation products are marked with asterisks. Intact peptide is denoted as M, and precursor ions are annotated as (M+G). All annotations of intact peptides retaining partial glycan correspond to Y fragments. Fragments matching neutral losses of water and –CO were not annotated.

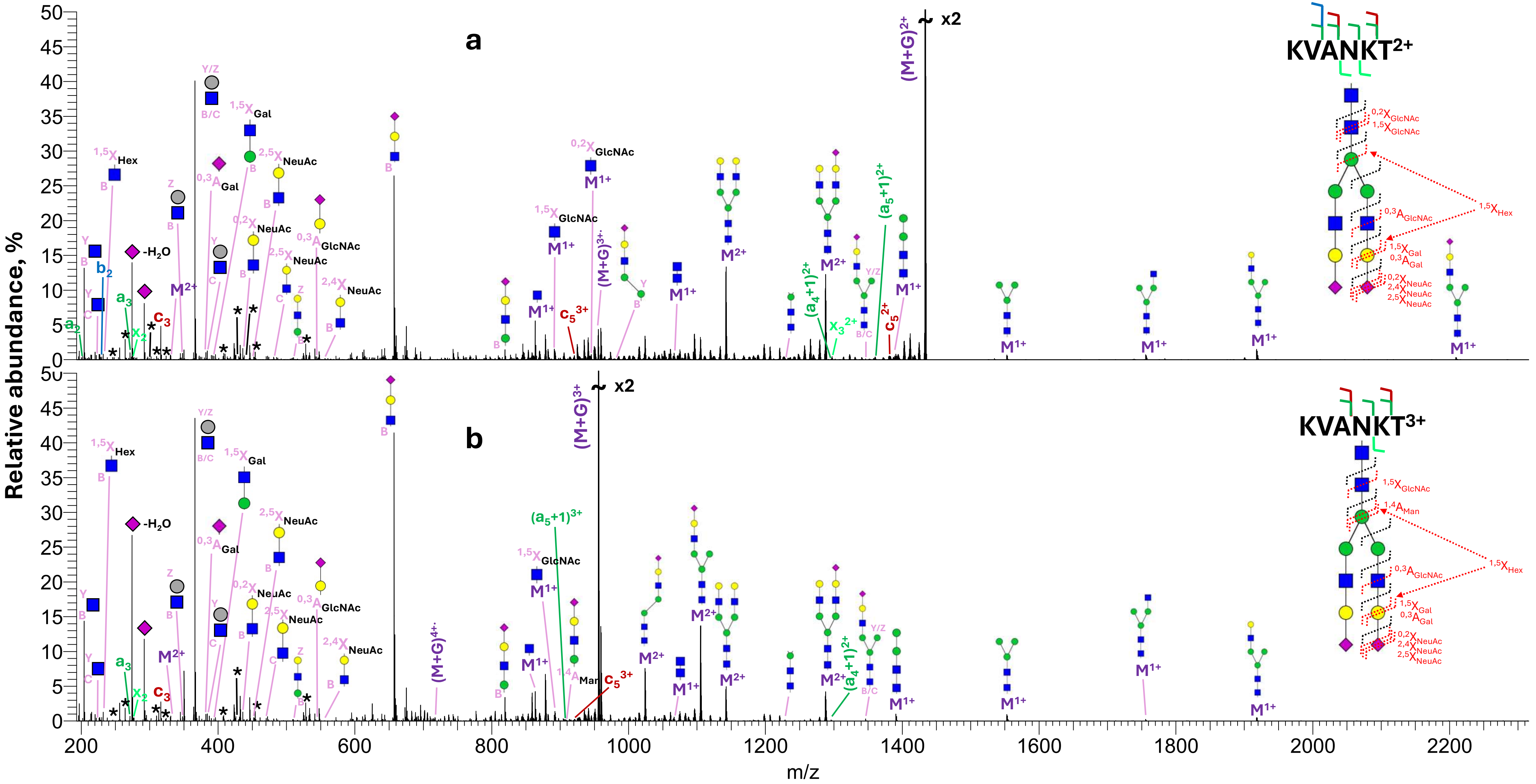


**Supplementary Figure S5**. Averaged MS2 EID mass spectra of doubly (**a**) and triply (**b**) charged A2G2S2 glycoform of the standard glycopeptide, acquired after irradiating the precursor ions for 150 ms (**a**) or 100 ms (**b**) by 25 eV electrons. M/z values matching to multiple isobaric internal glycan fragmentation products are marked with asterisks. Intact peptide is denoted as M, and precursor ions are annotated as (M+G). All annotations of intact peptides retaining partial glycan correspond to Y fragments unless otherwise specified. Fragments matching neutral losses of water were not labelled. Typically, only one charge state of a fragment was annotated.

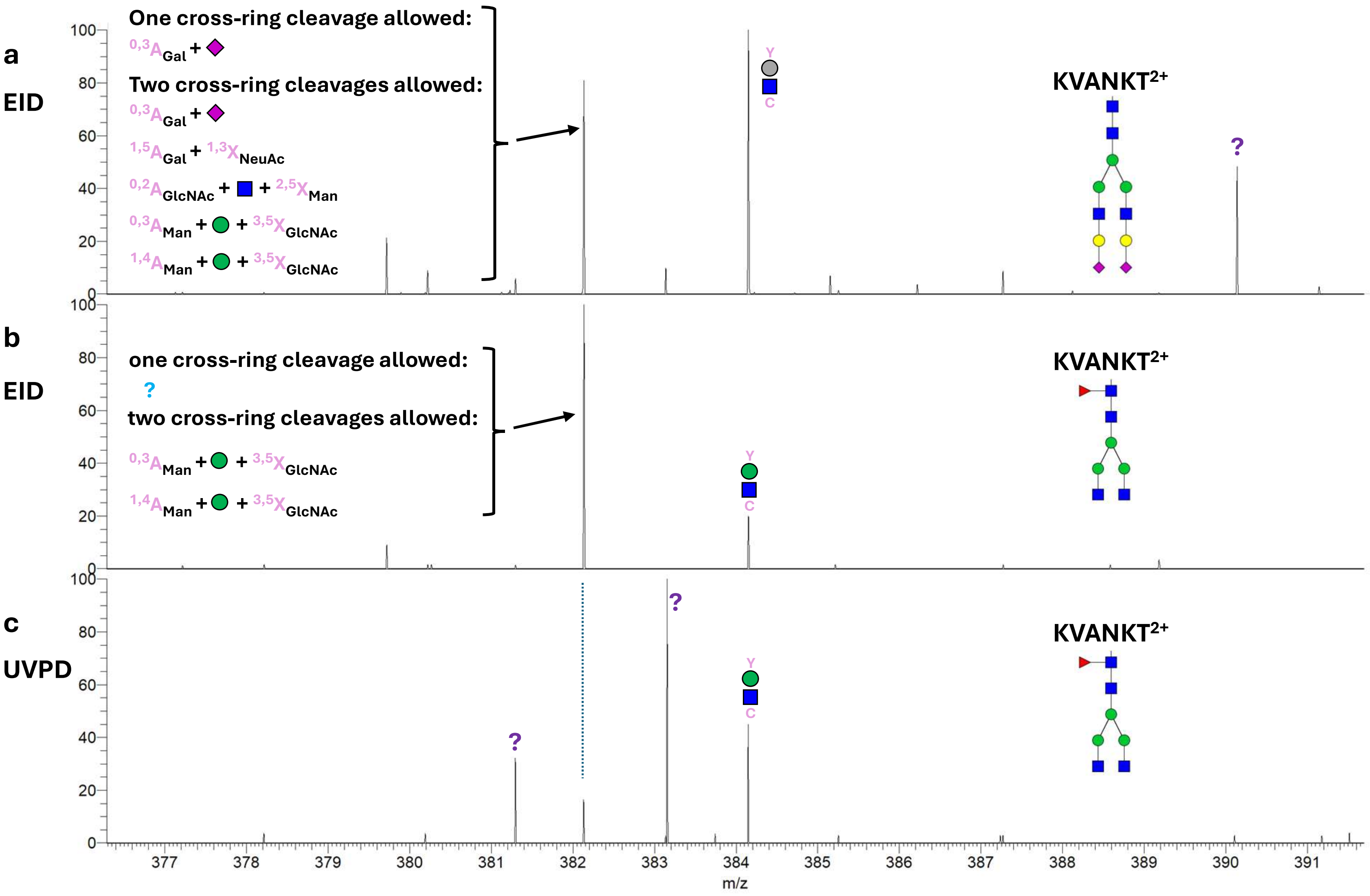


**Supplementary Figure S6**. Zoomed-in parts of mass spectra shown in Figures S6A **(a)**, S6B **(b)**, and S4B **(c)**, covering the m/z range between 377 and 391 Th. The fragment at m/z 382.134 may correspond either to $^{0,3}A_{Gal}$ fragment if only one cross-ring dissociation event is allowed or to multiple isobaric structures if two cross-ring dissociation events are allowed. All assignments are within 3 ppm mass accuracy.

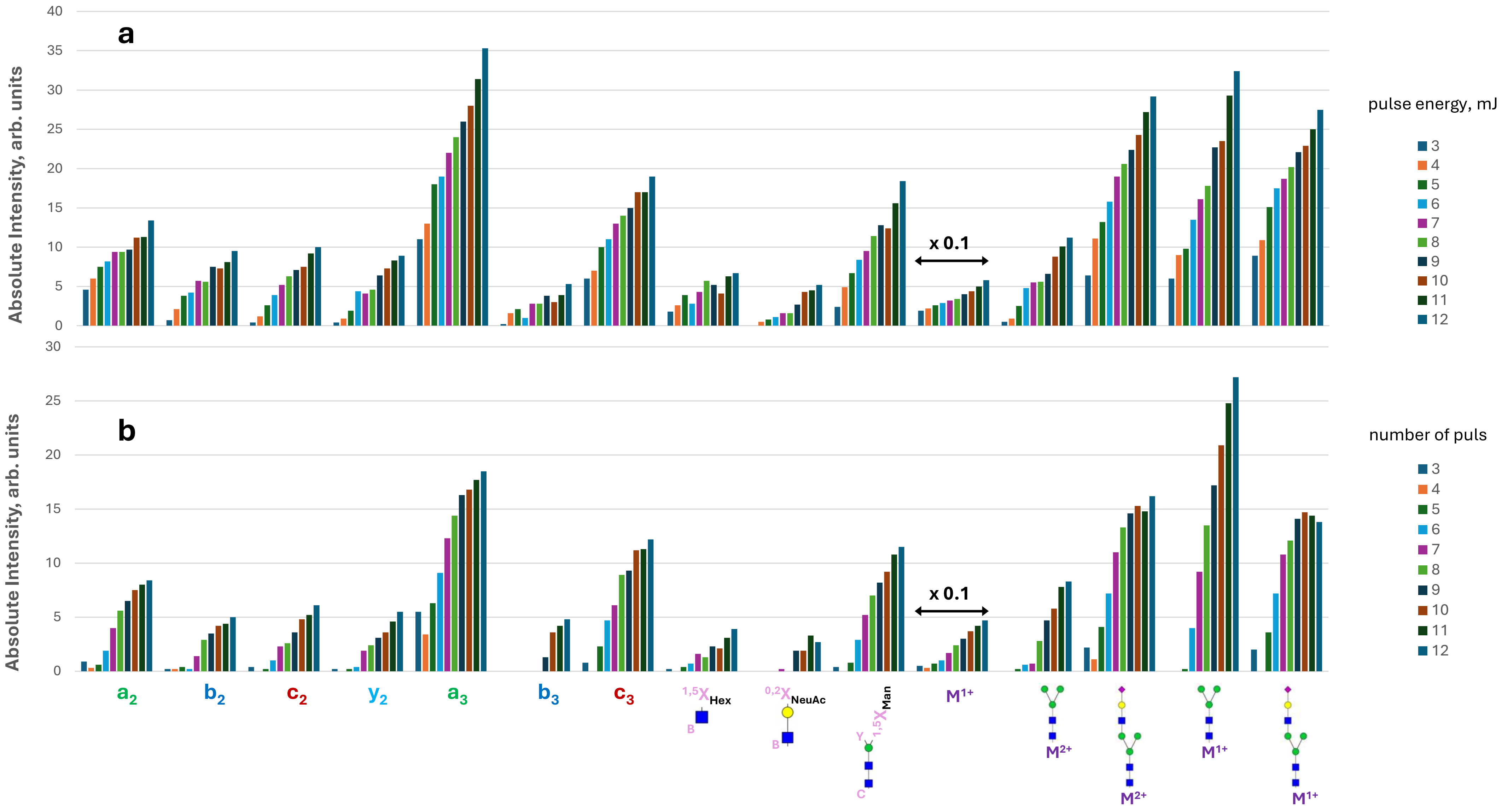


**Supplementary Figure S7**. Intensities of few selected fragments observed in averaged spectra acquired in direct infusion UVPD of doubly charged A2G2S2 glycoform of the standard glycopeptide using 8 laser pulses and different pulse energies (**a**) and different number of laser pulses at 10 mJ/pulse (**b**).

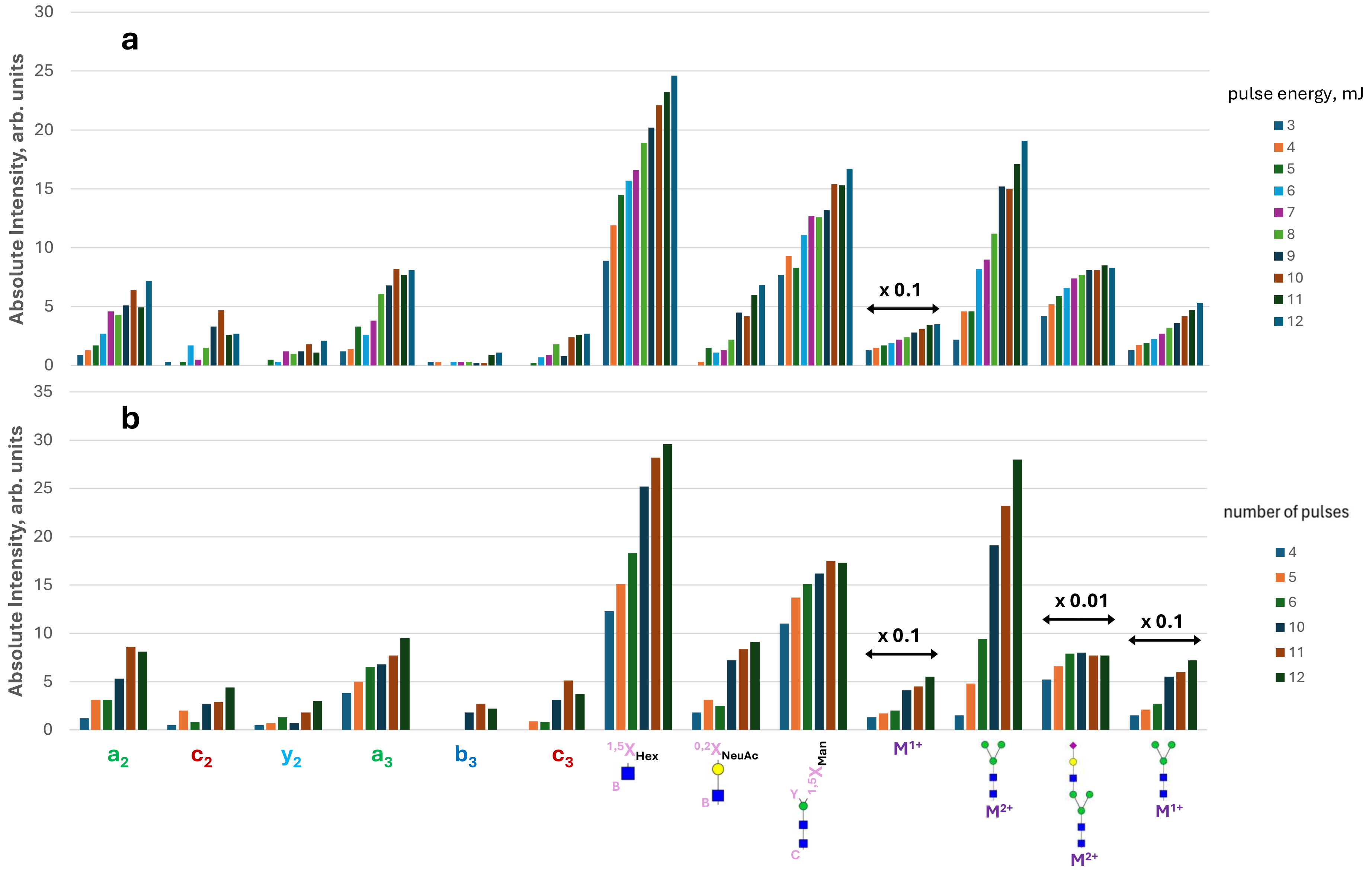


**Supplementary Figure S8**. Intensities of few selected fragments observed in averaged spectra acquired in direct infusion UVPD of triply charged A2G2S2 glycoform of the standard glycopeptide using 8 laser pulses and different pulse energies (**a**) and different number of laser pulses at 10 mJ/pulse (**b**).

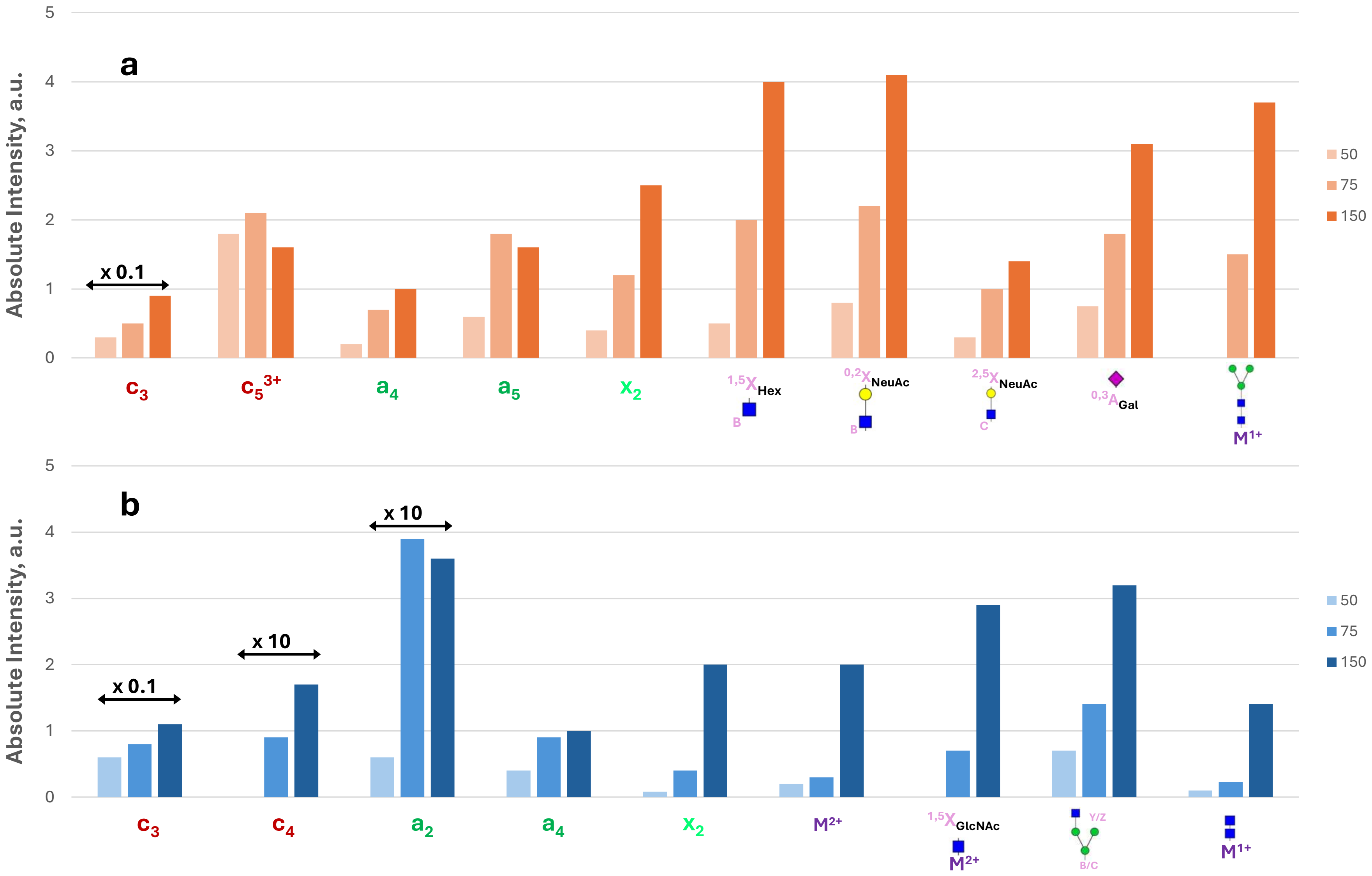


**Supplementary Figure S9**. Intensities of few selected fragments observed in averaged spectra acquired in direct infusion EID of triply charged A2G2S2 **(a)** and doubly charged FA2 **(b)** glycoforms of the standard glycopeptide using different irradiation times.

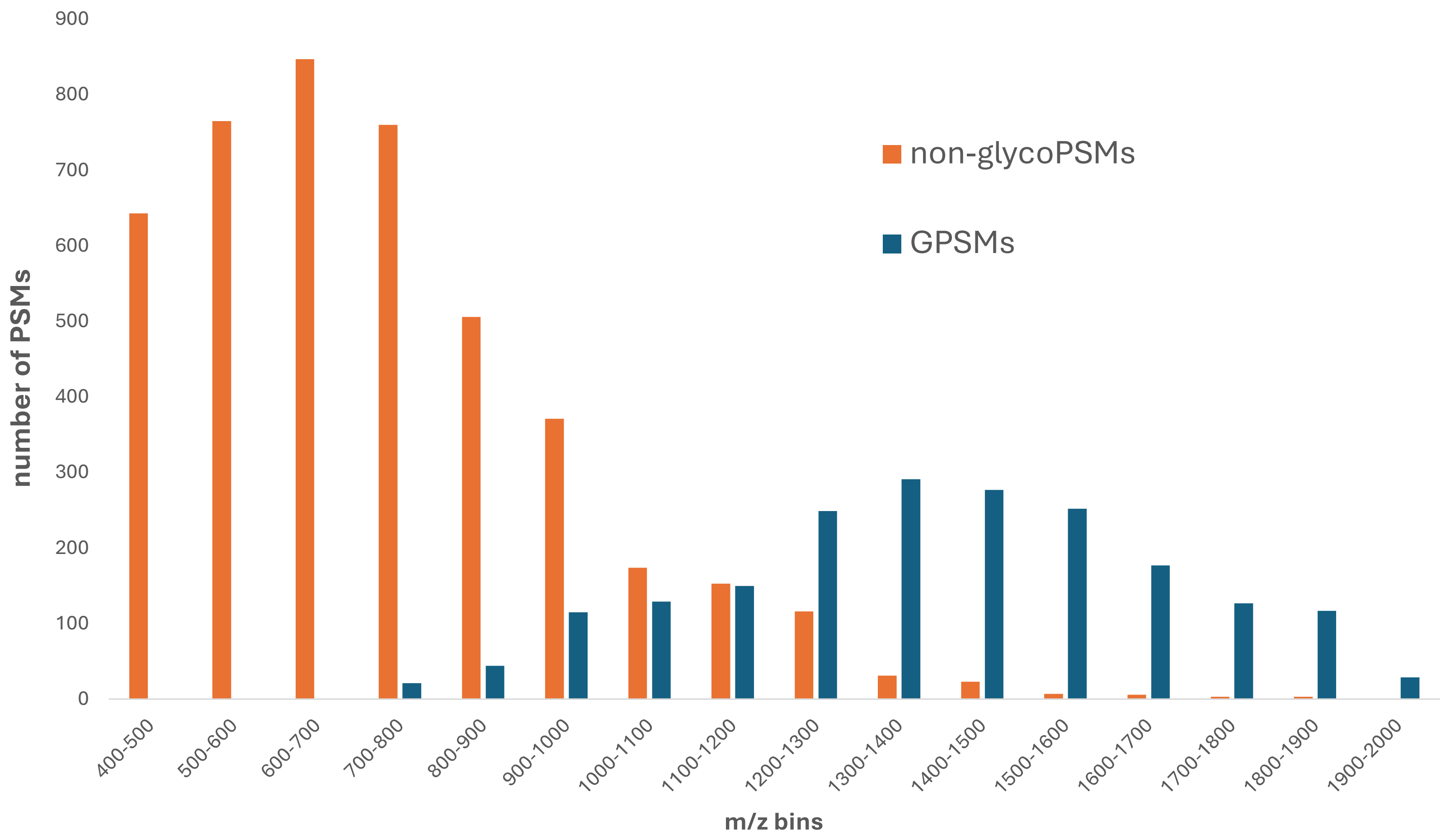


**Supplementary Figure S10**. M/z-binned distributions of the numbers of non-glycoPSMs (orange) and glycoPMSs (blue) identified in sceHCD analysis of complex glycopeptide mixture.

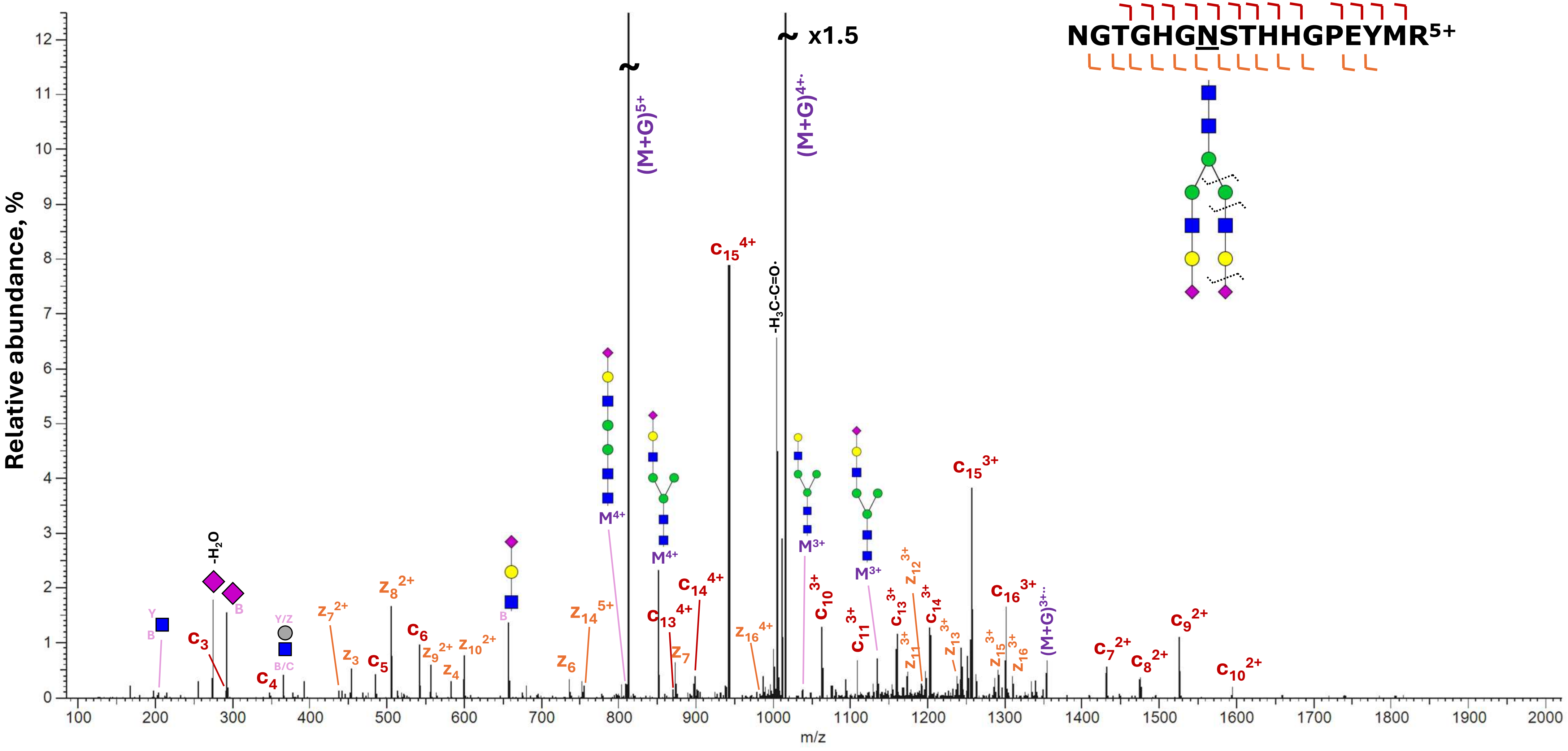


**Supplementary Figure S11**. ECD spectrum of N-glycosylated NGTGHGNSTHHGPEYMR$^{5+}$ acquired in LCMS of a complex glycopeptide mixture. Precursor ions were irradiated by electrons for 100 ms. Intact peptide is denoted as M, and precursor ions are annotated as (M+G). All annotations of intact peptides retaining partial glycan correspond to Y fragments.

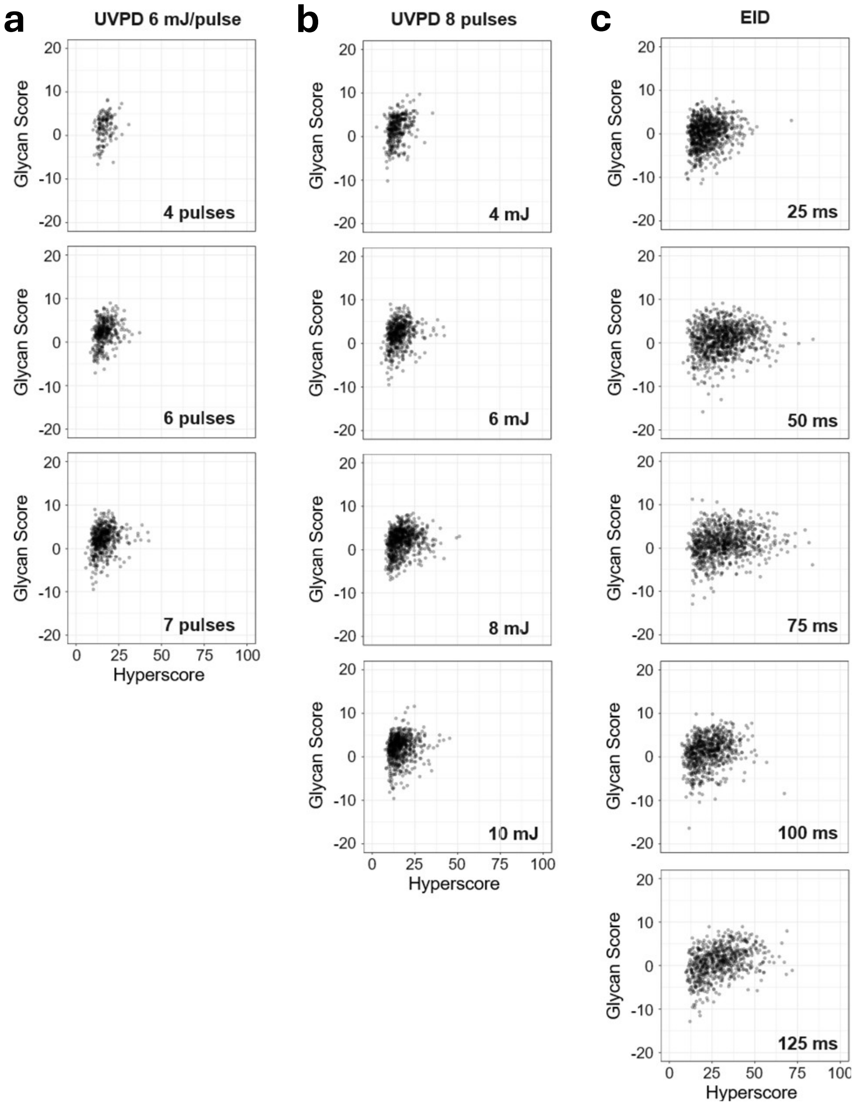


**Supplementary Figure S12.** Glycan scores plotted against hyperscores for glycoPSMs identified in UVPD using different numbers of laser pulses at 6 mJ/pulse (**a**), 8pulses at various pulse energies (**b**) and in EID experiments in a range of electron irradiation times (**c**). *B,y* types of ions were used in UVPD data analysis, and *b,y,c,z* peptide fragments were used in in the analysis of EID data.

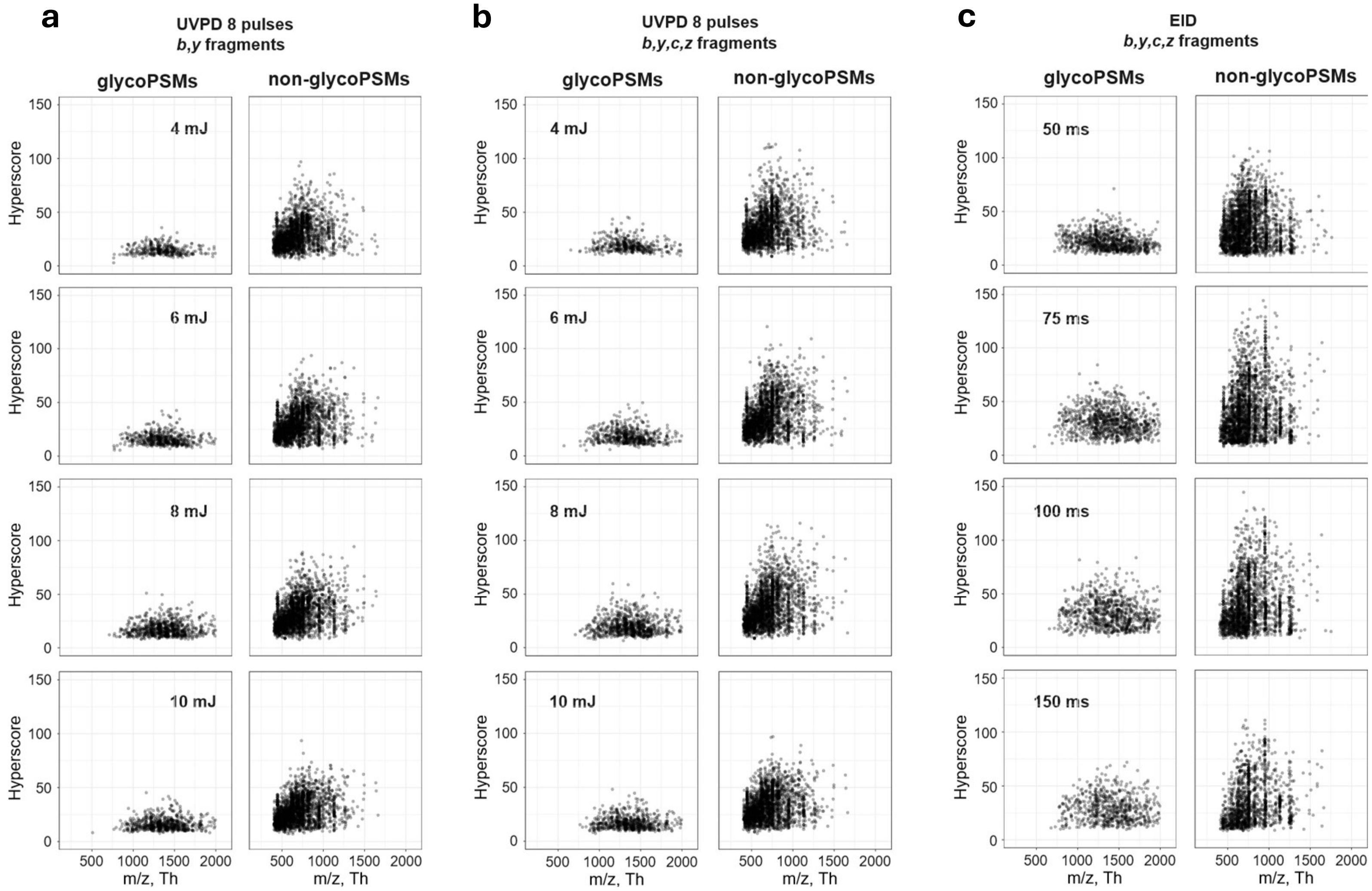


**Supplementary Figure S13.** Hyperscores plotted against *m/z* of glycoPSMs and non-glycoPSMs identified in UVPD using 8pulses at various pulse energies and *b,y* fragments for analysis (**a**), 8pulses at various pulse energies and *b,y,c,z* fragments for analysis (**b**) and in EID experiments in a range of electron irradiation times and *b,y,c,z* fragments for analysis (**c**).
